# Liquid-liquid phase separation (LLPS) as a sensing and adaptation mechanism: An evidence-based hypothesis on AP2 transcription factors in the malaria parasite

**DOI:** 10.64898/2025.12.10.693351

**Authors:** Valentín Iglesias, Yunuen Avalos-Padilla, Oriol Bárcenas, Xavier Fernàndez-Busquets

## Abstract

**Background:** Protein liquid-liquid phase separation (LLPS) can be driven by prion-like domains (PrLDs) inside intrinsically disordered regions (IDRs). The causing agent of the deadliest form of human malaria, *Plasmodium falciparum*, has abundant prion-like proteins whose aggregation is presumed to have a functional role. Multiple members of the largest family of transcription factors in *P. falciparum*, AP2 (*Pf*AP2), responsible for adapting the parasite gene response in different scenarios, were found in aggregation-prone protein screenings.

**Results:** We show that the *Pf*AP2s members carry the physicochemical determinants to perform LLPS forming biomolecular condensates *in vivo*. The long IDRs of *Pf*AP2s could sense changes in the cellular microenvironment, and their PrLDs could drive conformational rearrangements. *Pf*AP2s do not function as centralizing hubs for protein-protein interaction networks, but display significant preferred interactions among themselves, establishing a large, connected subnetwork. Predictions suggest that all *Pf*AP2s have regions to localize into LLPS-condensates, while larger *Pf*AP2s could initiate condensation. We show that four *Pf*AP2 members co-localize in live *P. falciparum*, bearing the potential to be engaged in LLPS-condensates.

**Conclusion:** We present bioinformatics analyses and experimental data obtained in live parasites suggesting that *Pf*AP2s are able to direct LLPS in *P. falciparum*. We propose a model in which sensing by the parasite of cellular stresses like host transfer, temperature changes and energy depletion, and the corresponding gene responses are driven by LLPS where the *Pf*AP2 family plays a fundamental role. Finally, we postulate targeting *Pf*AP2 as a new therapeutic antimalarial strategy to curb the emergence of drug-resistant parasites.

## Introduction

The concept that not all proteins require a precisely defined three-dimensional (3D) structure to execute their biological functions underscores the remarkable diversity and adaptability within the field of protein functionality. While traditional views have often emphasized the importance of well-folded structures for protein function, emerging research has unveiled a myriad of proteins that exhibit functional versatility without conforming to a rigid 3D framework. This flexibility is particularly evident among intrinsically disordered proteins (IDPs) and intrinsically disordered regions (IDRs) within proteins. These regions lack a fixed tertiary structure yet play crucial roles in various cellular processes, including signalling, regulation, and molecular recognition [1, 2]. Instead of adopting a single, static conformation, IDPs and IDRs have a dynamic flexibility to interact with diverse binding partners, enabling them to participate in intricate cellular networks and pathways [3]. IDPs/IDRs populate a continuum of degrees in their content of order and disorder, ranging from compact to fully extended conformations [3]. This plasticity is highly dependent on physicochemical conditions and binding partners, being in most cases necessary to develop their functions and a common mechanism to regulate them [1]. Yeast prions are modular proteins [4, 5] which contain a subclass of IDRs named prion domains (PrDs), typically enriched in asparagine (Asn) and glutamine (Gln) [6–8]. PrDs are capable of existing in at least two conformations: a soluble form and usually a self-propagating aggregate [8, 9]. The change in conformation is commonly accepted as a beneficial trait, a bet-hedging strategy that costs individual yeast cells to grow slower but in turn allows populations to thrive in suboptimal conditions [10–12]. Proteins displaying prion-like characteristics have been termed prion-like proteins and are mostly identified by having low compositional complexity (LCC) domains analogous to yeast prions, which are referred to as prion-like domains (PrLDs) [13]. Both prions and prion-like proteins are largely involved in the regulation of gene expression through modification of the affinity of complexes that bind DNA, DNA compaction, or RNA processing [14–19].

PrLDs can form diverse types of membraneless structures by themselves or by interacting with nucleic acids [20–25]. These are commonly referred to as biomolecular condensates as their different functions usually stem from their capacity to concentrate specific molecules in a defined space [20, 26]. Biomolecular condensates are thought to be formed by the spontaneous separation of multivalent proteins that stably coexist in two states, a dilute and a dense phase, in a process known as liquid-liquid phase separation (LLPS) [27, 28]. In this process PrLDs would act as regulators of the highly dynamic phase transitions involved in the formation and dissociation of these condensates. The interactions sustaining phase separated condensates depend heavily on the physicochemical parameters of the system and are therefore tightly regulated through pH, temperature, salt concentration or post-translational modifications among others [22, 23, 29, 30]. Mutations, post-translational modifications, peaks of protein concentration or decline in their solubility can cause a transition towards aberrant assemblies, which usually entails the deposition of amyloid fibrils [20, 22, 31]. Such transitions have been related to different disorders in humans including neurodegeneration and cancer [20, 22]. Not only liquid to solid transitions can be deleterious to cells, but also the impairment of molecule exchange between the condensate and the dilute phases or the abolishment of condensate formation or dissociation can lead to pathological conditions in cells or cell populations [32, 33]. LLPS as a mechanism for the organization of subcellular compartments is a conserved strategy already seen throughout diverse phyla with examples in bacteria [34], fungi [35–37], plants [24, 38], animals [27, 39], and even in plastids [40–42] and virus [43, 44].

Condensates regulate gene expression [45, 46] in the cytoplasm (e.g. RNA maturation) or the nucleus (e.g. mediator complex), and have been proposed to act as a fast response to changing environmental conditions, usually stress [37, 47–49]. Transcription factors (TF) typically consist of globular DNA-binding domains and LCC IDR activation domains [46, 50], which regulate gene transcription establishing protein-protein interactions (PPI) with coactivator complexes and other TFs [50, 51]. Recent discoveries identified various TFs developing their highly specific expression programs via condensate formation [36, 46, 52–57].

Malaria is a tropical disease caused by parasites of the genus *Plasmodium*. The World Health Organization estimated for 2023 263 million cases and 597,000 deaths, which confirms the lack of progress in reducing both indicators since 2020, due in part to the emergence of strains resistant to currently used therapeutics [58]. To reverse this trend and achieve malaria eradication, novel antiplasmodial strategies will be needed, anticipating resistance emergence by understanding how the parasite relates with the changing surrounding environment. Among the human-infecting species, *Plasmodium falciparum* is the parasite responsible from 85% of cases and most casualties annually [58]. The *P. falciparum* genome has over 80% AT bases [59] and the resulting UT-rich transcripts show preference towards encoding Asn to the point where an estimate of 30 % of the parasite’s proteins contain LCC IDRs, especially poly-Asn or Asn-rich domains [60]. First estimates indicated that an astonishing one-fourth of the *P. falciparum* proteome could have prion-like propensity [61], which was more recently lowered to ca. 10% of its proteins adjusting for the parasite’s biological background [62]. Moreover, we showed that these PrLDs have amyloid cores capable of forming amyloid-fibrils [62].

In *Plasmodium*, only one major family of TFs has been identified, the apetala2 (AP2), composed of 28 proteins [63]. This is a surprisingly low number compared to over 1600 TFs in humans [51] and 2296 in *Arabidopsis thaliana* [64]. AP2 TFs are found in plants and Apicomplexa (the phylum to which *Plasmodium* belongs) but absent from other eukaryotes like mammals or yeasts [63]. The malaria parasite AP2 family contains one to four AP2 DNA-binding domains of approximately 60 residues, and, similarly to their plant homologues, each domain forms a triple-stranded beta-strand stabilized by an alpha-helix structure [63, 65]. As in plants, these short AP2 domains tend to be the principal globular element in the protein [63]. Whereas in *A. thaliana* AP2 TFs are mostly small proteins around 300 amino acids [66], *P. falciparum* AP2s (*Pf*AP2s) range in length from 200 to 4109 residues (**Figure 1**). Interestingly, while the globular AP2 domains are essential for modulating DNA-binding, these proteins remain largely unstructured. In *P. falciparum* the 28 proteins with AP2 DNA-binding domains are expressed throughout the life cycle, where they mediate transcriptional regulation of genes and chromatin condensation specific to the different parasite stages [67].

**Figure 1 –.**
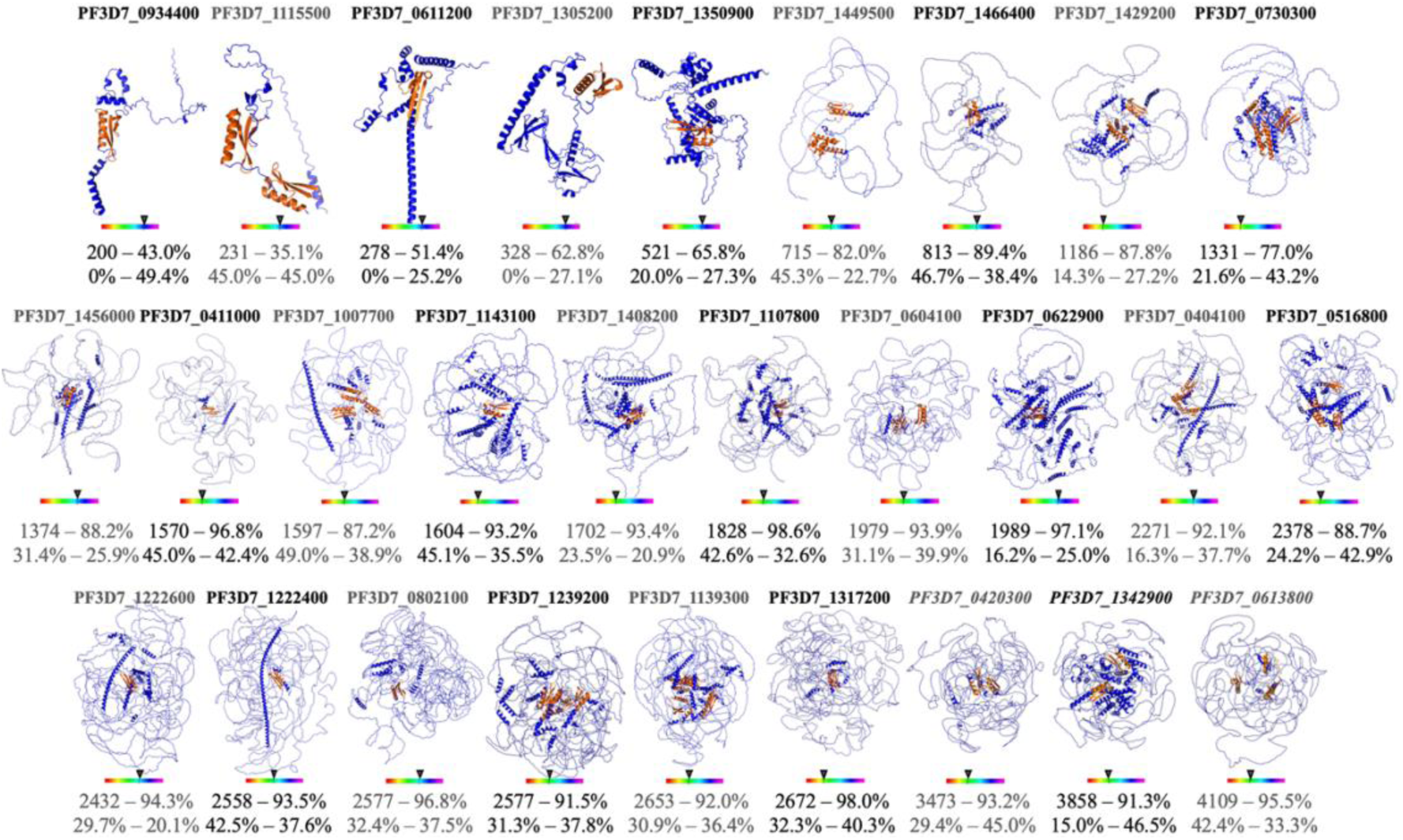
Characterization of the identified *Pf*AP2 protein family. Computational structural models for the *Pf*AP2s in blue except the globular AP2 domains which are depicted in orange. The protein isoelectric point (pI) is outlined as a pH one-dimensional plot ranging from pH 1 (coloured red) to pH 14 (coloured violet). The first line below the structure lists the protein length and the percentage of disordered residues according to AlphaFold pLDDT. The second line corresponds to the percentage of residues predicted to undergo binding-dependent and pH-dependent conditional ordering, respectively. *Pf*AP2s structures were extracted from the AlphaFold database except those missing (*Pf*ApiAP2 PF3D7_0420300, *Pf*AP2-HS PF3D7_1342900 and *Pf*ApiAP2 PF3D7_0613800, indicated in italics), for which models covering the globular AP2 DNA-binding domains are shown (regions 1600-3473, 2000-3858 and 2400-4109, respectively).

*Pf*AP2s regulate gene expression by directly binding DNA through their AP2 globular domains. They can functionally homodimerize as seen in the crystal structure of *Pf*AP2-SP/EXP [65], heterodimerize with other members of the family like *Pf*AP2-G and *Pf*AP2-I [68], and interact with a variety of nuclear proteins [68–73] or with chromatin [68, 70, 72, 74, 75]. A recent study by Shang et al. identified at least nine AP2 members that could form such a chromatin-regulatory complex [75], and a subsequent work by Subudhi et al. co-precipitated a chromatin remodelling complex with 6 different AP2s [76]. Protein stretches which mediate interaction often overlap with aggregation-prone regions [77] even in non-hydrophobic polar IDRs [78]. We previously identified protein aggregation in all *P. falciparum* stages [79] and showed that PfAP2s self-association could become aberrant as we identified at least nine AP2 members that formed aggregates in late blood stages (*Pf*AP2-O, *Pf*AP2-SP/EXP, *Pf*AP2-O5, *Pf*APiAp2 PF3D7_0613800, *Pf*AP2-11A/P, *Pf*AP2-O2, *Pf-*SIP2, *Pf*ApiAP2 PF3D7_0420300) and demonstrated that different peptides from the IDRs of *Pf*AP2-HS were capable of forming amyloid fibrils [79, 80].

This combined background led us to the hypothesis that stress-sensing and gene expression in *P. falciparum* may be driven by LLPS mechanisms, of which *Pf*AP2 TF members would be cornerstones. If this was the case, destabilization of *Pf*AP2-driven condensates could be an interesting and yet unexplored target for future antimalarial approaches. Here, as a starting point we computationally explored the potential of *Pf*AP2 TFs to undergo LLPS with the help of state-of-the-art bioinformatics tools. Among all members of the family, we selected four *Pf*AP2 candidates and experimentally validated their co-localization inside *P. falciparum* cells further stressing their potential to modulate the parasite gene expression via phase separated intermolecular assemblies.

## Results

### PfAP2 are mostly disordered, large proteins

Recent developments in protein structure prediction allowed us to structurally characterize the *Pf*AP2 TF family, since there are only four solved structures available; two of them covering the globular AP2 domains [65], and the other two depict solely the folded AP2-coincident C-terminal domain (ACDC) [81], which represents a small portion of the proteins. We obtained model structures for 25 *Pf*AP2 from the AlphaFold database and generated protein models for the remaining three *Pf*AP2 (see **Methods**) [82, 83]. The computational structural models align with previous sequence-based predictions [63], as these are mostly disordered proteins, with AP2 DNA-binding domains being the main sometimes the only folded element in the TF (**Figure 1** and **Figure S1**). Notably, the AP2 globular domains in AlphaFold structures resemble in all cases those in the experimentally resolved structures for *Pf*AP2-SP/EXP and SIP2 (Protein Data Bank (PDB) IDs: 3IGM and 6SY0) [65]. AlphaFold models report a local confidence metric known as the predicted local distance difference test (pLDDT) which confers higher values to correct local structure. Remarkably, low pLDDTs has shown to be a strong reporter of disordered or conditionally disordered domains [83–86]. Analysis of the *Pf*AP2s revealed that they are mostly IDPs, and their disorder/order rate increases with their length (p-value < 0.001).

Intriguingly, state-of-the-art disorder prediction methods failed to detect a significant degree of disordered regions in the *Pf*AP2s (**Table S1**) [87, 88]. As previously pointed out, a significant degree of this cryptic disorder could correspond to partially folded or conditionally folded domains [86, 89]. We explored ANCHOR2 [87] which reports energetically compatible regions able to perform disorder-to-order transition upon binding and DispHred [90], which applies pH-dependent hydrophobicity and charge to discriminate between ordered and disordered regions (**Figure 1**). For most of the *Pf*AP2 members, disorder is predicted to be dependent on the binding partners (88%, 22/25), except for 3 of the 4 shortest proteins. For protein environmental pH, all *Pf*AP2s showed a significant degree of pH-conditional folding (**Figure 1**). These results suggest that many of these IDRs could mediate conditional folding by adopting particular structures in the presence of specific binding partners (e.g. large macromolecules or metabolites) or host conditions that entail pH changes in the cellular environment. Conditional folding is not uncommon for IDPs, which can acquire different conformational states when interacting with specific binding partners or functional assemblies [91]. In particular, the IDRs of nuclear receptor coactivators CBP and ACTR’s undergo mutual synergistic folding upon interaction [92], while upon binding the transcriptional activator JUN, PAGE4 undergoes disorder-to-order transition [93]. In humans, IDPs with prion-like domains have been shown to sense pH variations, where TFs could modify gene expression upon slight (<1) pH shifts [94]. Additionally, pH fluctuations sufficed to control the phase-separation behaviour of human RNA-processing proteins [95]. In a similar way, disorder-to-order transition of IDRs in *Pf*AP2s could help to sense changes in the surrounding pH, e.g. as a secondary messenger for energetic starvation or as a sign of changing hosts (e.g. transition from human circulation to mosquito and vice versa), to stabilize the DNA-binding capacity of the folded AP2 domains or to interact with different proteins.

### PfAP2 proteins are highly interconnected

Proteins rarely carry out their biological roles independently; instead, they require diverse inputs to be functional. For *Pf*AP2s, functional interactions between them [65, 68] or with different *Plasmodium* proteins have been characterized [68–72]. We measured the connectivity of the *Pf*AP2 family as a proxy of their cooperativity by applying associations from the STRING (Search Tool for Retrieval of Interacting Genes/Proteins) database [96]. STRING bypasses the lack of proteome-wide experimental evidence for physical binary annotation by also including data from multiple sources: organism-specific databases, predictions from homologues, text-mining results, genomic content and also shared pathways, all in all providing a high proteomic coverage. The final evidence is weighted in a score that favours data obtained from multiple sources rather than from a single site. To restrict as much as possible low-confidence interactions or those coming from a unique source we applied a high confidence cutoff (>0.700), a threshold for which 32.9% of the *P. falciparum* proteome (1769 proteins) lacked confident associations. Under these conditions 27/28 *Pf*AP2 (except the short *Pf*AP2 PF3D7_1115500 protein) had annotated PPI.

Singh and Gupta estimated the number of uncharacterized proteins in different databases for *P. falciparum* to be around 30% [97], and although this lack of confident annotation is especially manifested for understudied organisms, even human interactome maps are deemed largely incomplete [98]. This lack of annotation is clearly a bottleneck for different bioinformatics approaches and may degenerate in an unintended annotation bias between well-studied against less-studied genes [98]. As the *Pf*AP2s have been previously studied in different contexts and to avoid a potential bias, we discarded computing all the *Pf*AP2s against the whole *P. falciparum* and instead compared to those genes for which data is available. In this scenario, the 27 *Pf*AP2s have an average of 38.70 interactions against 36.72 of the interacting background (**Figure 2A**), which dispels the idea of the *Pf*AP2s functioning as hubs of PPI. Focusing on the interactions that *Pf*AP2 proteins carry out among themselves revealed that these are unequivocally higher than those expected by chance (**Figure 2B**) (*Pf*AP2s 80 interactions, random 7.5; p-value < 10^-5^). Finally, we explored the distances of the *Pf*AP2s in the network and compared them to those randomly expected by calculating two complementary measures: the size of the largest connected component and the mean shortest distance [98]. The first parameter is the largest connected subgraph and the second corresponds to the shortest distance of each *Pf*AP2 to other *Pf*AP2 (**Table 1**). Despite the incompleteness of the interactome, measurements of the mean shortest distances suggest that *Pf*AP2 proteins are agglomerated in the same neighbourhood of the interactome network (scores of 1 correspond to agglomerated proteins and 3 to completely scattered), and the largest connected component describes a degree of 25 interconnected *Pf*AP2 proteins. The overall network results show clearly that *Pf*AP2s share a highly interconnected interactomic vicinity, larger than randomly expected.

**Figure 2 –.**
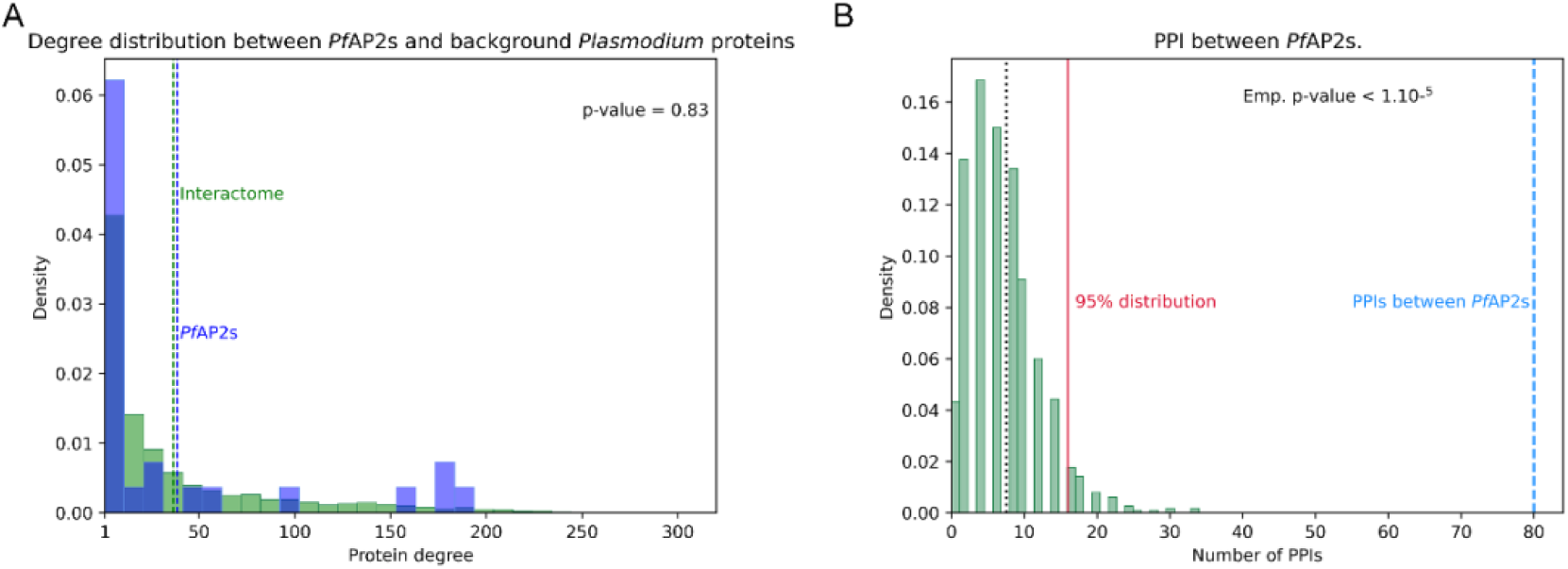
*Pf*AP2 interactome. **A)** Degree distribution for the association-annotated proteins in *P. falciparum* (n=3607, coloured green) and the *Pf*AP2s with annotated PPIs (n=27, coloured blue). **B)** Number of PPIs between *Pf*AP2s (n=27; dashed blue line) compared to 1000 sets of random sampling from the proteins with described PPIs (median shown as dotted black line, 95% distribution as a solid red line).

**Table 1.**
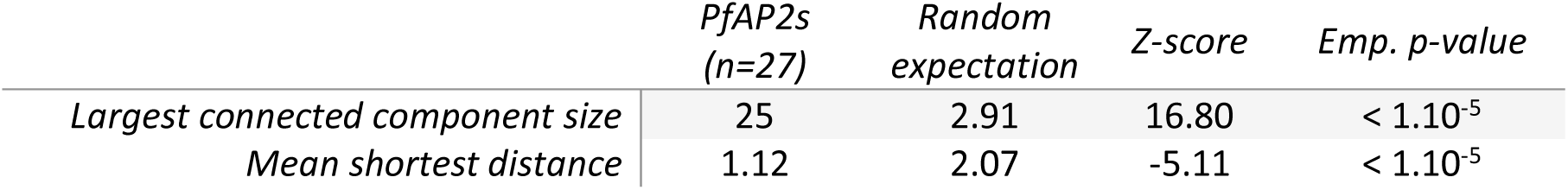
*Pf*AP2 proteins are located closer in the network than randomly expected.

|  | *PfAP2*s<br>(n=27) | Random<br>expectation | Z-score | Emp. p-value |
| --- | --- | --- | --- | --- |
| Largest connected component size | 25 | 2.91 | 16.80 | < 1.10<sup>-5</sup> |
| Mean shortest distance | 1.12 | 2.07 | -5.11 | < 1.10<sup>-5</sup> |

### Most members of the PfAP2 family bear predicted PrLDs

PrLDs resemble yeast prion domains by having LCC (especially biased in Asn and Gln residues) which promote structural disorder conformations. The presence of prion-like proteins and their role in modulating gene translation as nucleic acid binding elements are widely conserved among phyla [9, 16, 19, 62, 99, 100]. As different structural conformers from several prion or prion-like proteins can be passed through cellular divisions and alter the gene expression patterns in absence of changes to the genetic sequence, these proteins can act as epigenetic elements [11, 101]. In a recent study, through a highly stringent approach, we characterised the *P. falciparum* prionome and showed that one regulator of gene expression, the translation initiator factor IF2c, had inside its PrLD an amyloid core capable of aggregating into fibrillar amyloid assemblies [62]. Analysing the prion-like potency of the *Pf*AP2 TF family with the PLAAC prediction method revealed that most members (n=18) contain a predicted PrLD (**Table 3** and **Figure 3**). The distribution of PrLDs shows a distinction between shorter proteins where no *Pf*AP2 under 700 residues has a predicted PrLD, while 17 out of the longest 22 proteins (77.3%) are predicted to have at least one of these modular regions. These 18 identified PrLDs have amyloid potential as in all cases pWALTZ detects amyloid cores inside them (**Table S2**) [102].

**Figure 3 –.**
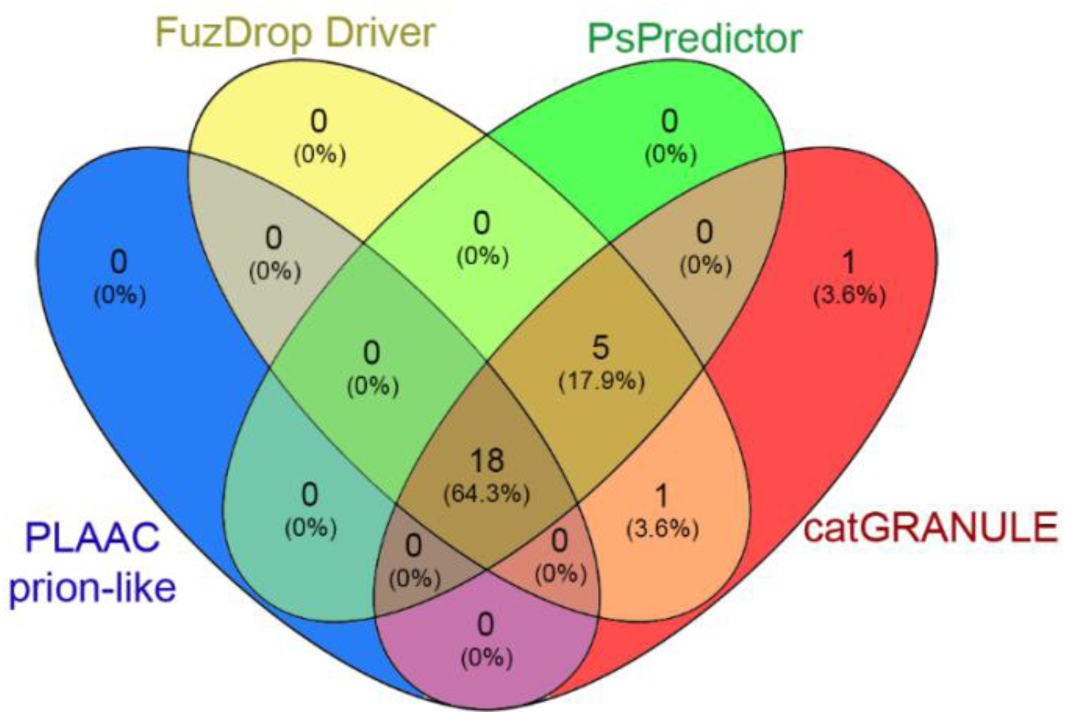
Phase separation and prion-like predictions convergence on *Pf*AP2s. Venn diagram showing the concordance of predictions for the *Pf*AP2s by LLPS prediction methods FuzDrop, PSPredictor, catGRANULE and PLAAC prion-like prediction. The number of positive hits are indicated and, in parentheses the percentage they represent relative to the whole *Pf*AP2 family (n=28).

### Bioinformatics tools predict that PfAP2s form biomolecular condensates

Proteins with IDR PrLDs are usually found forming or recruited to different classes of condensates, in many cases necessary and sufficient for driving phase transitions [103]. The current stickers and spacers model for LLPS proposes that sticker regions contribute the attractive multivalence interactions while spacers provide sufficient flexibility to withstand phase separation [104]. Folded domains, interaction motifs or individual residues can act as stickers. Different studies carried out in the FUS protein family have reported the key role played by Tyr and Arg residues as stickers and how their content and distribution can influence the phase behaviour of PrLDs, determining a solid or liquid-like state of the compartment [103–107]. Tyr residues were also necessary for insulin receptor coalescence and signalling [52]. In multiple proteins mutating Tyr impaired or attenuated LLPS; [24, 52, 53, 107, 108], whereas the content of Arg in dipeptides from the protein C9orf72 correlated with their LLPS potency [109]. Notably, in the human hnRNPDL protein, only the isoform which includes both the Tyr- and Arg-rich IDRs could phase separate spontaneously [107]. Tyr has been reported to mediate hydrophobic and π-stacking interactions, while Arg residues can form electrostatic interactions or hydrogen bonds. Together, Arg and Tyr are capable of establishing cation-π interactions [30, 109]. We therefore analysed if the sequences of the *Pf*AP2s would bear Tyr- or Arg-rich domains using fLPS’s single-residue bias calculation (**Table 3**) [110]. Notably, 26 members (92.9%) carried a Tyr-rich domain with the only exception of two of the shortest proteins (*Pf*ApiAP2 PF3D7_1115500 and *Pf*AP2-SP/EXP). This was not the case for Arg, as only 10 *Pf*AP2s (35.7%) have an Arg-rich region. This result points out that not all members would be able to phase separate based on cation-π interactions, but a large part of them could through hydrophobic and π-stacking.

Proteins which drive the formation of the condensate are usually referred to as scaffolds or drivers, while proteins which are recruited are called clients [111]. In general scaffolds tend to have higher valences than clients, thus being able to establish more interactions [112], and are compositionally biased towards disorder promoting residues (Pro, Gly, Ser) and prion-forming residues (Asn and Gln), and depleted in order-promoting residues (Phe, Iso, Val, Cys, Trp) [112]. To further characterize the capacity of the *Pf*AP2 TF family to spontaneously drive phase separation, we applied recently developed software: catGRANULE, FuzDrop and PSPredictor [113, 114]. catGRANULE estimates the propensity of proteins to form condensates by weighting nucleic acid binding, protein disorder and in a less extent the residue bias towards Phe, Gly and Arg and protein length [115]. PSPredictor is a highly accurate machine-learning prediction method based on the data from the LLPS-database and especially capable of detecting proteins drivers of LLPS [114, 116]. DeePhase combines knowledge-based features and a custom neural-network based language model to identify LLPS-driving proteins especially trained for distinguishing between LLPS and non-LLPS IDPs [117]. FuzDrop is based on the observation that, in order to drive LLPS multivalent and non-specific interactions, conformational entropy should be driving these condensations [112]. This tool is capable of distinguishing entropy-driven droplet-promotion, aggregation and binding from input sequences and sorting out drivers of LLPS from clients [112, 113]. These last three algorithms were among the top-scoring in a recent benchmark for drivers of LLPS [118]. Analysis on the capacity of *Pf*AP2s to scaffold LLPS showed a similar pattern (**Table 3** and **Figure 3**), whereby all members larger than 700 residues were classified as drivers and only three *Pf*AP2s were not predicted as positives. catGRANULE includes as putative positive hits the 328 residue *Pf*ApiAP2 PF3D7_1305200 and the 521-residue *Pf*AP2-O4, the latter also scoring slightly above the FuzDrop driver threshold. FuzDrop predicts all the shorter *Pf*AP2 members to bear the determinants to be incorporated into pre-formed condensates. Notably, FuzDrop assigns 22% of *P. falciparum*’s proteome as capable of driving LLPS (**Table 3 and Table S3**) a lower number than previous estimates for the human proteome [112]. Resampling this proteomic result by 1000 random datasets of equal size to the *Pf*AP2s (n=28) revealed that the potential of *Pf*AP2s to drive LLPS was significantly higher than randomly expected (p-value < 1.10^-5^).

### In vivo co-localization of LLPS-predicted PfAP2s

Highest confidence LLPS drivers (according to the previously mentioned prediction methods) are proteins whose length (over 2000 and up to >4000 residues) greatly impairs exogenous expression and purification, running into solubility issues, aggregation, or difficulty in achieving a significant amount of protein. To overcome these problems, we selected as a proof-of-concept *Pf*AP2-O5, one of the smallest *Pf*AP2 predicted to form condensates with high-confidence. However, *Pf*AP2-O5 recombinant expression in different *Escherichia coli* bacterial systems resulted in very low yields (**Figure S2**). Because of these hurdles and the impossibility to work directly with the most likely *Pf*AP2 LLPS scaffolds in *P. falciparum*, we decided to take advantage of their ability to bind specific DNA sequences [67, 75] allowing to indirectly target these proteins and decipher their possible condensate formation *in vivo*.

We selected four *Pf*AP2s expressed at different timepoints throughout the parasite asexual blood stages considering: their predicted role as driver of LLPS (**Table 3** and **Figure 4**), their expression patterns covering all parasitic asexual blood stages, and divergency in their DNA sequence recognition. For *Pf*AP2-I, *Pf*ApiAP2 PF3D7_0420300, *Pf*AP2-SP/Exp and *Pf*AP2-12, oligonucleotides composed of their target DNA sequence were synthesised with different fluorophores capable of performing Förster resonance energy transfer (FRET) when in close proximity (**Table 2**).

**Figure 4 –.**
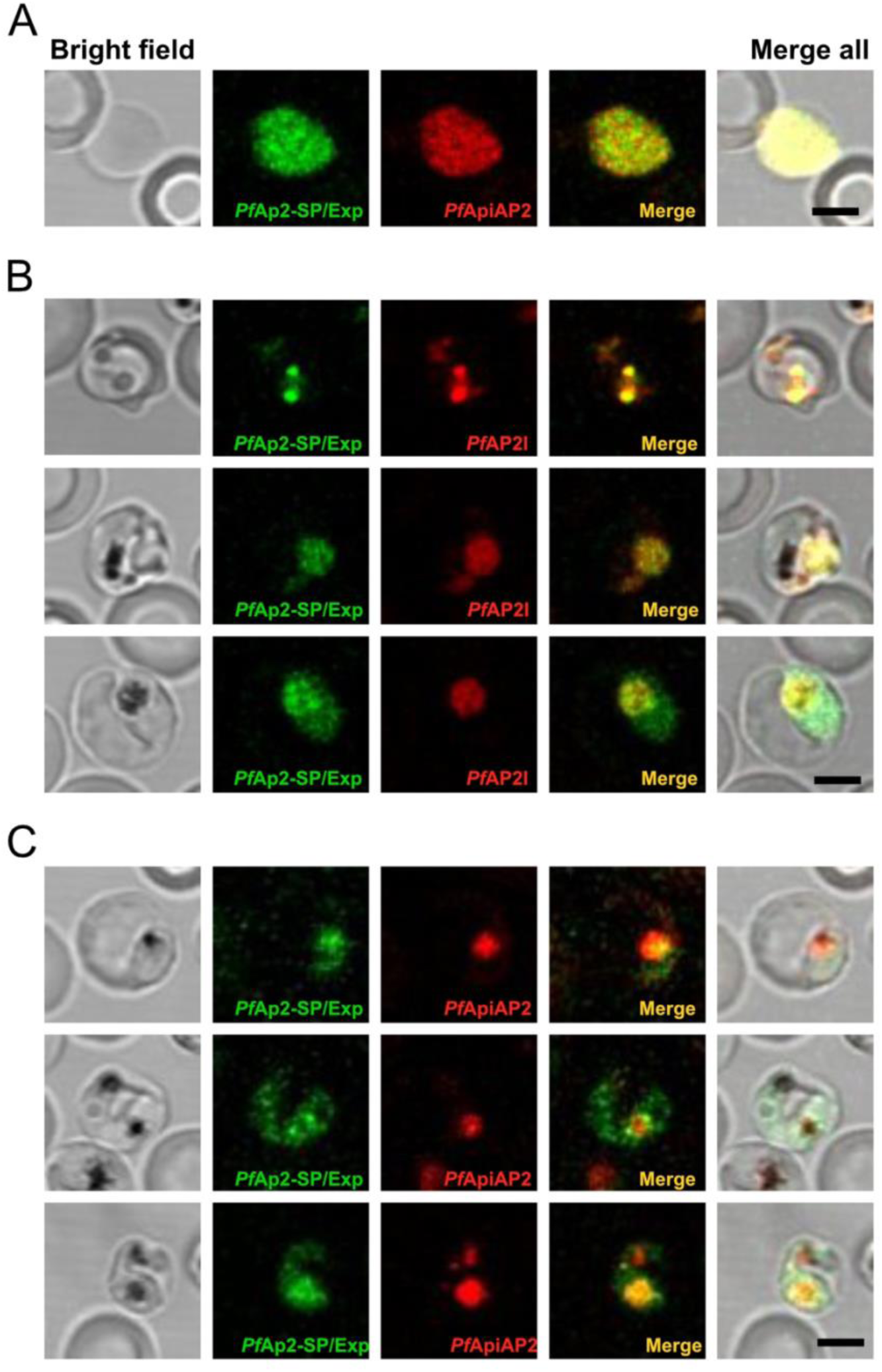
Confocal fluorescence study of the *Pf*AP2s distribution inside live *P. falciparum*-infected ghost red blood cells (gRBC). Red blood cells (RBCs) were initially subjected to hypotonic lysis, and fluorescently labelled oligonucleotides present in the medium entered gRBCs before resealing. These gRBCs were infected with *P. falciparum* parasites, and those parasitized gRBCs (pgRBCs) containing both oligonucleotides were imaged by confocal fluorescence microscopy. Oligonucleotides targeting *Pf*AP2-SP/Exp (tagged with Alexa 488 fluorophore) are shown in green colour, and those targeting *Pf*AP2-I and *Pf*ApiAP2 PF3D7_0420300 (tagged with Alexa 546 fluorophore) are shown in red colour. **A)** Non-parasitized gRBC containing both oligonucleotides. **B, C)** Examples of three pgRBCs in the presence of oligonucleotides targeting **B)** *Pf*AP2-SP/Exp and *Pf*AP2-I, and **C)** *Pf*AP2-SP/Exp and *Pf*ApiAP2. Scale bar = 5 μm.

**Table 2.** *Pf*AP2 proteins and their specific DNA-binding sequences selected for FRET analysis.

| *PfAP2* | **AP2-I** | **ApiAP2** | **AP2-SP/Exp** | **AP2-12** |
| --- | --- | --- | --- | --- |
| AP2-recognized dsDNA sequence | #5'GGTGCACTA<sup>3'</sup> | #5'GTGTTACAC<sup>3'</sup> | *5'TGCATGCA<sup>3'</sup> | *5'TCTACAAA<sup>3'</sup> |

5'-end fluorescent tags: # Alexa 546 (acceptor), \* Alexa 488 (donor)

A modified version of the lyse-reseal erythrocytes for transfection (LyRET) protocol [80, 119] was followed to load tagged oligonucleotides into reconstituted or “ghost” red blood cells (gRBCs) prior to *P. falciparum* infection. Briefly, erythrocytes were incubated in a hypotonic lysis buffer in the presence of oligonucleotides to allow their entry during transient membrane permeabilization and before cell resealing. This procedure had a negligible effect on *P. falciparum* growth compared to the standard parasite culture using intact erythrocytes) [80]. Confocal fluorescence microscopy imagining revealed the entry of both oligonucleotides in correctly lyse-resealed gRBCs (**Figure 4A**). In the presence of *P. falciparum*, the homogenously distributed fluorescence of both oligonucleotides that was characteristic of non-parasitized gRBCs was observed to coalesce into discrete regions inside the pathogen (**Figure 4B, C**). Notably, puncta-like structures become visible in a significant number of parasitized gRBCs (pgRBCs), a common structural arrangement seen in different membrane-less condensates.

FRET efficiencies in oligonucleotide-loaded gRBCs were significantly different between non-parasitized and parasitized. In non-parasitized gRBCs, FRET signals were negligible (0.01-0.2%), while in pgRBCs, FRET efficiency depended on the donor/acceptor combination, ranging from 10 to 70% **(Figure 5)**. These FRET results indicated that the selected *Pf*AP2 pairs were located within nanometer scale proximity in the parasite cytoplasm, consistent with distances typical of biomolecular condensates. Notably, FRET efficiencies for *Pf*AP2-SP/EXP as donor were higher with *Pf*AP2-I and *Pf*ApiAP2 than having as donor *Pf*AP2-12. It is tempting to speculate that this difference could be a reflection of the existence of different kinds of condensates formed by *Pf*AP2s, although further co-localization experiments with other *Pf*AP2s should be performed to confirm this point.

**Figure 5 –.**
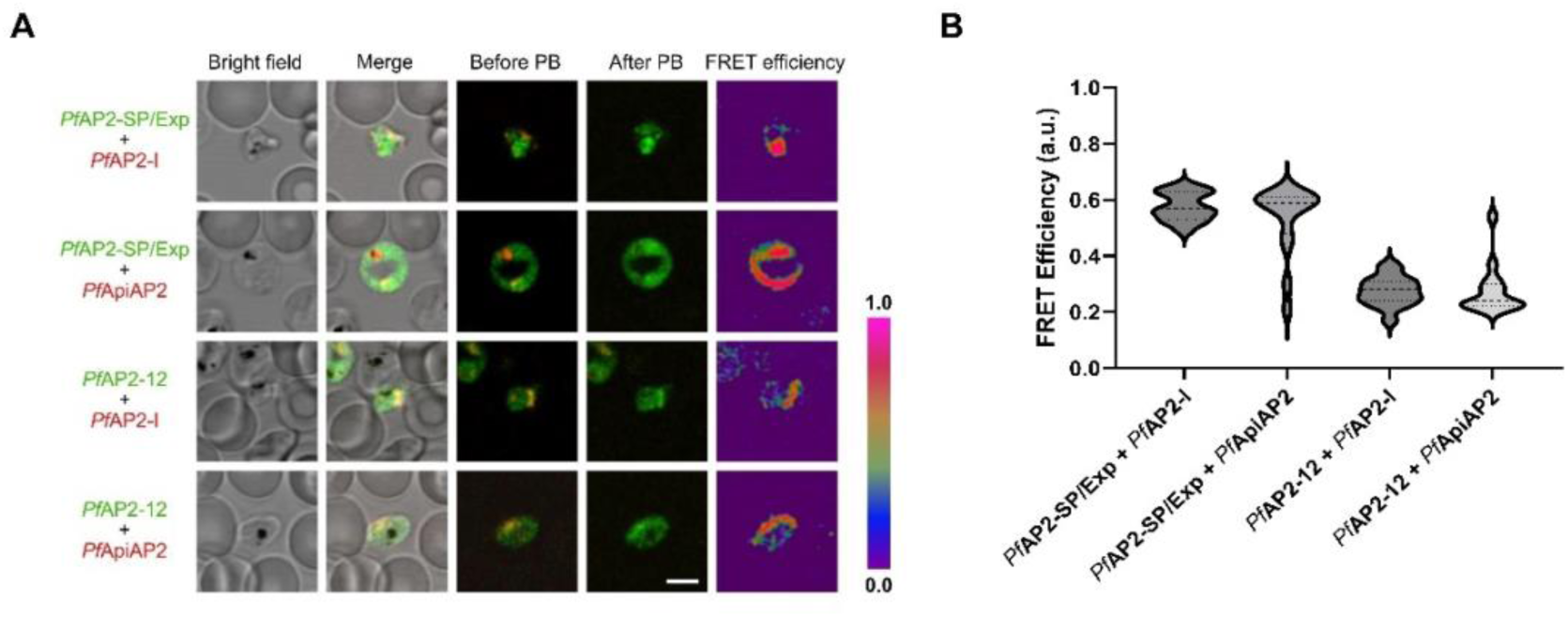
Indirect reporters of *Pf*AP2 co-localization in live *P. falciparum* cells. **A)** Representative images showing co-localization of *Pf*AP2 pairs in *P. falciparum-*parasitized gRBCs, identified by the presence of hemozoin crystals. **B)** FRET efficiency calculated for the different *Pf*AP2 pairs. Oligonucleotides targeting *Pf*AP2-SP/Exp and *Pf*AP2-12 acted as donors (tagged with Alexa 488 fluorophore, green colour), and those targeting *Pf*AP2-I and *Pf*ApiAP2 PF3D7_0420300 acted as acceptors (tagged with Alexa 546 fluorophore, red colour). Scale bar = 5 μm. PB, photobleaching. a.u, arbitrary units.

## Discussion

IDPs/IDRs have moved from a neglected position following the structure-function paradigm to the realisation of their critical roles in binding, regulation or scaffolding [1–3]. Their difficulty to crystalize meant they were predominantly absent from 3D structures and their identification was mostly carried out through sequence-dependent prediction methods [120]. AlphaFold structural models for the malaria parasite’s biggest transcription factor family, *Pf*AP2 reflects large disordered proteins where the DNA-binding AP2 domains are in most cases the only structural element [82, 83]. Classical disorder prediction methods fail to classify the totality of disorder in *Pf*AP2 proteins depicted in AlphaFold models. Previous works suggested that this discordance could be reflecting the transientness of some secondary structure; including the potential of an IDR to fold upon different biophysical conditions [86, 121]. Conditional folding in IDPs can be driven by binding to other IDPs or globular proteins, temperature, pH and redox changes, or recognition of different sequential elements [78, 122–125]. The human splicing factor U2AF is an essential heterodimeric protein member of the spliceosome. The U2AF large subunit causes an induced fit on the unstructured small monomer via its N-terminal IDR [126] and, in a similar way, to an IDR from the splicing factor SF1 through its C-terminal [127]. Additionally, the U2AF large domain drives the LLPS of nuclear speckles with pyrimidine-rich RNA sequences [128]. Intracellular pH variations can modify the DNA binding affinities of TFs by protonating His residues, resulting in regulation of gene expression [94]. In a similar way, Lys, Asp (but also Glu and Arg) residues can upshift and downshift their pKas cellular ranges of 7.0-7.8 depending on their protein and electrostatic environments [129, 130]. The protonation of acid Asp and Glu residues from c-Myc PEST region (201-268) induces a disorder to order transition [131] and Prothymosin ɑ, which is unstructured at neutral pH, gains a compact, ɑ-helical rich conformation at low pHs as its negative charges get neutralized [132]. We explored the potential of stretches inside the *Pf*AP2s to perform inducible folding by means of binding partners or changes in pH. Our data suggest that a significant percentage of the *Pf*AP2 disordered domains could undergo pH- or binding partner-dependent disorder-to-order transition. Therefore, and in addition to their large sizes, these proteins could mediate the intermolecular interactions required for the transcription regulation but also physically ‘sense’ different environmental conditions. Parasites lacking a functional *Pf*AP2-HS, the master regulator of heat-shock response in *P. falciparum* [69, 133], showed severe growth defects already at 37 °C, and an increased sensitivity to proteostatic toxicity induced by treatment with artemisinin, the current frontline antimalarial drug [134]. Modifications in messenger molecules or temperature and pH variations when changing hosts from endothermal mammal to ectothermic mosquito could also affect *Pf*AP2 dynamics or their protein-protein interactions and instantaneously adapt the gene expression to succeed in these new situations. For example, *Plasmodium* gametocytes undergo an almost immediate molecular cascade towards gametes differentiation upon being exposed to an abrupt decrease in temperature combined with an increase in pH and xanthurenic acid [135].

The scarcity of TFs in *Plasmodium* suggests that their role in controlling gene transcription could be less redundant than for other eukaryotes. Previous attempts to knockout or knockdown members of the *Pf*AP2 family have had diverse outcomes. Most efforts to silence the *Pf*AP2s acting mainly as TFs were impracticable, leading to either unviable organisms or parasites unable to progress past a certain stage [71, 133, 136]. Notably, a truncated version of *Pf*AP2-SP/EXP which only expressed the globular AP2 domain could rescue the parasite viability in blood stages, although their target genes were markedly deregulated and therefore, causing the parasites to lose the immune evasion advantages that the antigenic variation provides [137]. On the other hand, the *Pf*AP2s associated with regulating heterochromatin condensation were viable and showed no clear signs of disfunction [75]. These have been predicted to form complexes with other AP2s, chromatin and chromatin-modifying proteins, with some members possibly having partially overlapping functions. For instance, together with *Pf*AP2-Tel, telomerase integrity and length maintenance are thought to be mediated (at least partially) by *Pf*AP2-G2, *Pf*AP2-G5 or *Pf*AP2-12 [75]. Considering their characteristics, their low redundancy and crucial role, disrupting their function could not only be tackled by disrupting a structural architecture but also their ability to perform PPI. The *Pf*AP2 interaction network represents a group of proteins that, far from being a hub centralizing a myriad of PPI, relies instead on establishing multiple interactions among themselves, forming a highly connected subgraph (**Figure 2** and **Table 1**). This result adds up to the hypothesis of *Pf*AP2s exerting their biological roles in close contact, possibly through biomolecular condensate formation.

A large number of *Pf*AP2s, and especially those larger than 700 residues, have predicted PrLDs capable of phase-transition in response to changing conditions. Prion-like proteins can act as a sensors in response to fluctuations in environmental conditions by a self-propagating change in conformation, either revealing hidden genetic load or as a fast-response to change in environmental conditions [15–17]. Such rapid response has been identified in multiple organisms of diverse phyla and is mostly associated to the regulation of gene expression [9], and therefore assumed to be an evolutionary conserved trait [138]. The yeast translation terminator factor protein Sup35 forms aggregated prions that allow read-through stop codons, revealing a previously hidden genetic load which provides individuals with a selective advantage in adenine deficient environments [139, 140]. Additionally, the bacterial prion and master translation terminator factor Rho is regulated by nucleotide levels, being kept as an aggregate in limited resource conditions but able to form soluble functional complexes when ATP/ADP intracellular levels rise [141]. Rho in the aggregated prion state allows bacteria to explore transcriptional variations, which are beneficial under stress conditions [16]. Nonetheless, such analogic adaptation only allows the system to be in two states and as the endpoint is usually an amyloid-like aggregate, reverting to the initial state is highly energetically demanding. Prion-like proteins can also mediate the formation of liquid condensates, mainly by establishing a multitude of low-specific interactions throughout their PrLDs [21, 23, 24, 27]. Accordingly, the formation of condensates offers multiple advantages to cells as an adaptation to fluctuating environmental conditions, allowing rapid responses that require only conformational changes in already synthesised proteins. This dynamic mechanism is amenable to regulation as the size, location, diversity and concentration of its components can be appropriately modulated [38, 142]. The human heat shock factor1 (HSF1) condensates at the heat-shock-protein gene loci to promote the transcription of heat-shock response genes [143]. These condensates are disassembled by chaperones, attenuating transcription upon cessation of thermal stress. In yeast, the Ded1p helicase reversibly phase-separates through its IDRs in response to heat and pH, repressing housekeeping mRNAs, promoting stress response and arresting cell growth [35]. In *Arabidopsis*, the prion-like protein FLOE1 functions as a hydric sensor [144]. In seeds FLOE1’s localization is diffuse in dry conditions and phase separates upon hydration, delaying germination until hydric conditions are favourable. Different plant IDPs and prion-like proteins condense in response to temperature, enhancing the organism’s thermotolerance. The prion-like protein FUST1 forms condensates through its PrLD that promote stress granule formation, enhancing thermotolerance [145], while different heat-shock transcription factors (HSFs) undergo condensation to promote transcription of heat-shock response genes including chaperones [146].

Biomolecular condensate formation driven by LLPS is an emergent field that tries to explain and characterize the existence of membraneless compartments. Previous reports have gathered notable evidence that Tyr and Arg interactions could mediate the LLPS formation of condensates, especially in human proteins such as FUS or hnRNAP [103–106]. A search for Tyr-and Arg-rich regions in the *Pf*AP2 protein dataset revealed that while several proteins could perform these mostly through cation-π interactions, Tyr-rich regions were more abundant than Arg, to the point that 92.9% (26/28) of the members had a Tyr-rich stretch. Therefore, these stretches would be able not only to interact with Arg but to perform π-π interactions with themselves or with intra- or inter-molecular aromatic residues. Most members of the *Pf*AP2 family are predicted as drivers of LLPS by dedicated prediction methods (**Table 3**). catGRANULE predicts that 25 out of the 28 *Pf*AP2s could mediate nucleic acid co-condensation [115]. In addition, the LLPS-predictor FuzDrop identified droplet-forming domains for all the *Pf*AP2 members; and therefore, having the features necessary for being recruited to condensates as clients. Notably, all *Pf*AP2s larger than 700 residues are predicted by the assayed dedicated tools as capable of driving phase separation. Previous *in vivo* research in *E. coli* cells and *in silico* molecular dynamics simulations showed that the size of IDPs correlates with their LLPS capacity, with larger proteins capable of phase-separating at concentrations several orders of magnitude lower than their shorter variants [30, 147]. The extensive IDRs of the *Pf*AP2 members could therefore help the proteins condense even if required at low cellular concentrations. The pattern is similar to the one reported by the prion-like propensity prediction, in which no member under 700 residues had an identified PrLD, while 77.3 % of the larger did. Altogether, these results suggest a functional role for the large IDR domains present in the *Pf*AP2 proteins. Just as the DNA-binding domains of the *Pf*AP2s are sequence-specific, the length and composition of their IDRs could determine the proteins’ ability to condense and to recruit binding partners for different situations. This could help explain the lack of target gene regulation in observed in the mutant version of *Pf*AP2-SP/EXP lacking the IDRs [137], as well as the aberrant phenotype observed in a truncated version of the LLPS predictors-positive protein PTP7 lacking its LCC Asn-rich domain [148].

**Table 3.** Compendium of evidence that leads us to propose a LLPS-based mechanism for the function of the AP2 family of TFs in *P. falciparum*.

| Preferred Symbol | Gene ID | Uniprot Ac. | Protein Length | Intrinsic disorder (%) | catGRANULE score | FuzDrop score - role | PSPredictor score | DeePhase score | R/Y-bias | PLAAC PrLD |
| --- | --- | --- | --- | --- | --- | --- | --- | --- | --- | --- |
| *Api*AP2 | PF3D7_1115500 | Q8IIK9 | 231 | 35.1 | 0.72 | 0.15 – C | 0.06 | 0.32 | None | None |
| *Api*AP2 | PF3D7_0934400 | Q8I2H4 | 200 | 43.0 | 0.17 | 0.21 – C | 0.03 | 0.18 | R+Y | None |
| AP2-EXP2 | PF3D7_0611200 | C6KSV7 | 278 | 51.4 | 0.74 | 0.15 – C | 0.06 | 0.32 | Y | None |
| *Api*AP2 | PF3D7_1305200 | Q8IER4 | 328 | 62.8 | 1.43 | 0.20 – C | 0.46 | 0.39 | R+Y | None |
| AP2-O4 | PF3D7_1350900 | A0A5K1K8Z4 | 521 | 65.8 | 1.97 | 0.63 – D | 0.39 | 0.74 | R+Y | None |
| AP2-O5 | PF3D7_1449500 | Q8IKY0 | 715 | 82.0 | 2.77 | 0.98 – D | 0.93 | 0.78 | Y | PrLD |
| AP2-SP/EXP | PF3D7_1466400 | Q8IKH2 | 813 | 89.4 | 2.62 | >0.99 – D | 0.96 | 0.84 | None | PrLD |
| AP2-O3 | PF3D7_1429200 | Q8ILH2 | 1186 | 87.8 | 1.99 | 0.79 – D | 0.69 | 0.77 | Y | None |
| AP2-L | PF3D7_0730300 | Q8IBF6 | 1331 | 77.0 | 2.48 | 0.85 – D | 0.82 | 0.78 | Y | PrLD |
| AP2-HC | PF3D7_1456000 | Q8IKR9 | 1374 | 88.2 | 3.17 | 0.98 – D | 0.87 | 0.75 | Y | PrLD |
| AP2-Z | PF3D7_0411000 | Q9U0H8 | 1570 | 96.8 | 3.16 | >0.99 – D | 0.93 | 0.83 | Y | PrLD |
| AP2-I | PF3D7_1007700 | Q8IJW6 | 1597 | 87.2 | 2.61 | >0.99 – D | 0.95 | 0.78 | R+Y | PrLD |
| AP2-O | PF3D7_1143100 | Q8IHT5 | 1604 | 93.2 | 2.35 | 0.98 – D | 0.85 | 0.76 | Y | PrLD |
| AP2-G2 | PF3D7_1408200 | Q8IM14 | 1702 | 93.4 | 2.83 | >0.99 – D | 0.72 | 0.79 | Y | None |
| AP2-11A/P | PF3D7_1107800 | Q8IIS4 | 1828 | 98.6 | 3.12 | >0.99 – D | 0.95 | 0.81 | Y | PrLD |
| *SIP*2 | PF3D7_0604100 | C6KSN9 | 1979 | 93.9 | 2.77 | 0.94 – D | 0.94 | 0.77 | R+Y | PrLD |
| AP2-SP3/Tel | PF3D7_0622900 | C6KT65 | 1989 | 97.1 | 2.50 | 0.85 – D | 0.80 | 0.73 | Y | None |
| AP2-SP2 | PF3D7_0404100 | Q8I1Y8 | 2271 | 92.1 | 2.62 | 0.82 – D | 0.85 | 0.78 | R+Y | PrLD |
| AP2-O2 | PF3D7_0516800 | Q8I3U0 | 2378 | 88.7 | 2.98 | >0.99 – D | 0.89 | 0.76 | Y | PrLD |
| AP2-G | PF3D7_1222600 | Q8I5I9 | 2432 | 94.3 | 3.23 | >0.99 – D | 0.87 | 0.77 | Y | PrLD |
| *Api*AP2 | PF3D7_1222400 | Q8I5J1 | 2558 | 93.5 | 2.85 | 0.96 – D | 0.95 | 0.77 | Y | PrLD |
| AP2-LT | PF3D7_08<br>02100 | Q8IAM1 | 2577 | 96.8 | 3.38 | >0.99 – D | 0.96 | 0.76 | R+Y | PrLD |
| AP2-12 | PF3D7_12<br>39200 | Q8I531 | 2577 | 91.5 | 2.88 | >0.99 – D | 0.86 | 0.76 | Y | None |
| AP2-G5 | PF3D7_11<br>39300 | Q8IHX2 | 2653 | 92.0 | 2.87 | >0.99 – D | 0.89 | 0.77 | Y | None |
| AP2-G3 | PF3D7_13<br>17200 | Q8IEE4 | 2672 | 98.0 | 3.16 | 0.99 – D | 0.95 | 0.81 | R+Y | PrLD |
| ApiAP2 | PF3D7_04<br>20300 | Q8I1N6 | 3473 | 93.2 | 3.58 | >0.99 – D | 0.87 | 0.82 | R+Y | PrLD |
| AP2-HS | PF3D7_13<br>42900 | C0H5G5 | 3858 | 91.3 | 2.81 | 0.96 – D | 0.82 | 0.79 | Y | PrLD |
| ApiAP2 | PF3D7_06<br>13800 | C6KSY0 | 4109 | 95.5 | 3.16 | >0.99 – D | 0.87 | 0.78 | R+Y | PrLD |

Mounting evidence suggests that LLPS could be a property of multivalent proteins, for which different phase regimes and intermolecular interactions would define the experimental requirements for *in vitro* phase separation [149], a notion prevalent in the field of polymer chemistry [150]. Given our interest in replicating intracellular conditions in the malaria parasite, we focused on the specific sequence binding capacity of the *Pf*AP2 domains and selected four candidates with clearly distinct DNA target sequences and expression patterns throughout the *P. falciparum* blood stages. Confocal fluorescence microscopy imagining showed that while in non-parasitized gRBCs the fluorescence from the pair of oligonucleotides was homogenously distributed, in pgRBCs this signal partitioned into discrete regions inside the pathogen and, in a significant number of parasitized gRBCs, puncta-like structures could be identified. FRET reported close distances for the oligonucleotides targeting *Pf*AP2s in live *P. falciparum* cells, which are compatible with phase separated compartments. Of note, *Pf*AP2-12 as donor showed worse FRET efficiencies than *Pf*AP2-SP/EXP for *Pf*AP2-I and *Pf*ApiAP2 as acceptors. This difference could imply different kinds of condensates formed at different times or requiring different cellular conditions than those replicated in cell culture. *Pf*AP2-driven FRET interactions appeared to be mostly cytosolic, a localization of already been observed for *Pf*AP2-I [68] and *Pf*AP2-G, although in this last case its distribution varied depending on the parasite stage [151]. TFs are usually cytosolic until activated by different stimuli, when they translocate to the nucleus to regulate gene transcription [152]. Further experiments with other members of the *Pf*AP2 family should help to draw a more detailed scenario for their dynamic subcellular compartmentalization.

Artemisinin-derived therapies have helped to decrease the burden of malaria in the last decades, but the continuous emergence of resistant parasites has stalled this trend posing a risk on the vulnerable populations, specially pregnant women and children [153, 154]. As a counteroffensive, a significant effort is being done in the development of post-artemisinin treatments such as targeting pathogen-specific pathways or organelles (e.g., the apicoplast), drug repurposing, chemical adaptation, and the application of nanotechnological strategies such as encapsulation in targeted nanocarriers to reduce toxicity for the host and increase delivery to the pathogen [80, 155–157]. We believe that, if a significant degree of the parasite stress-sensing and gene regulation stems from the formation of the same macromolecular assemblies, impairing such a physiological master regulator would be a witty strategy that would in turn minimize the possibility of adaptation and resistance emergence. In addition, the lack of homologues in the human host could allow for a larger therapeutic window of *Pf*AP2-disruptor agents. Along this line, several members of the *Pf*AP2 family have been shown to be essential for parasite survival. This suggests a low redundancy in their gene expression regulation, further evidencing the difficulties that *Plasmodium* would face to overcome *Pf*AP2-targeted therapies. Recently, independent studies analysed potential strategies to target IDP-driven biomolecular condensates by means of small molecules [158–161], with promising results targeting the androgen receptor, a TF highly associated to prostate cancer [53] and the human respiratory syncytial virus replication [160]. A somehow evident approach could be to drive the pathogenic aggregation of specific, essential *Pf*AP2s or to accelerate a liquid to solid transition of the whole condensate causing a lack-of-function, but other strategies could imply to prevent their formation or disassemble those already preformed. These effects could be achieved by selecting peptides, aptamers or small molecules that accelerate or block those transitions. Mechanistically, these agents could stabilize an unbound conformation that keeps key proteins out of the condensate, weaken the interactions that maintain condensate integrity or stability, or reduce the free energy barrier required for pathogenic aggregation or condensate solidification. This scenario could be also valid for other human infecting parasites. Despite sequence variation, AP2 TFs in *Plasmodium malariae*, *Plasmodium knowlesi* and *Plasmodium vivax* maintain the physicochemical features that make predictive tools consider most of their members potential drivers of LLPS (**Figure 6** and **Table S5**). Other apicomplexans which are etiological agents of human tropical diseases such as babesiosis or toxoplasmosis also rely on AP2 for regulating gene transcription and could presumably be also targeted by such strategies.

**Figure 6 –.**
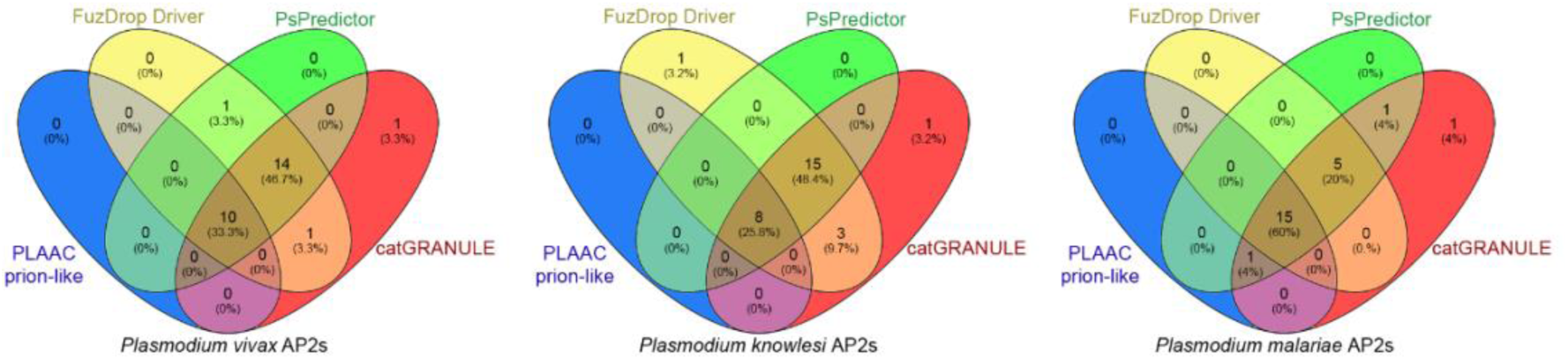
LLPS and prion-like propensity conservation in AP2s from *Plasmodium* species. Venn diagrams showing the concordance of the LLPS prediction methods FuzDrop, PSPredictor, catGRANULE and of PLAAC prion-like prediction for the AP2s of the human malaria-causing species: *Plasmodium vivax*, *Plasmodium knowlesi* and *Plasmodium malariae*. Three AP2s from *P. vivax* and *P. knowlesi* and two AP2s from *P. malariae* were not predicted as positives. The number of positive hits are indicated and, in parentheses, the percentage that they represent relative to the identified AP2 proteins (see **Methods** and **Table S5**; number of assayed proteins: 30, 31 and 25 respectively).

## Conclusions

We propose a model in which the sensing of different cellular stresses in *Plasmodium falciparum*, including host defences, temperature changes and energetic levels, and the corresponding gene expression responses are driven by LLPS in which the essential *Pf*AP2 family plays a fundamental role. We show that bioinformatics analyses predominantly point towards larger *Pf*AP2s being able to direct LLPS maintaining and adapting genetic responses. Subsequently, we show that different combinations of *Pf*AP2s reside in close proximity inside pRBCs, potentially forming LLPS-driven molecular condensates. Finally, we propose disrupting the *Pf*AP2 condensate formation as an invaluable antimalarial target for future therapies.

## Methods

### Disorder prediction

AlphaFold v2 [83] predictions were analysed for their pLDDT metrics and scores <70 were interpreted as disordered [83, 84]. LocalColabFold tool version v1.5.2 was used to model for proteins absent from the AlphaFold database (Q8I1N6, C0H5G5 and C6KSY0) [82]. MobiDB-lite disorder predictions were extracted from MobiDB [162]. ANCHOR2 and IUPred2a long disorder were extracted from IUPred2a web server [87]. DispHred pH-dependent disorder was obtained from the DispHred web server for the pH range of 3-9 and default settings (step of 0.5 and 51 residue window) [90].

### Protein-protein interaction analysis

The *Pf*AP2 family was scanned for associations with data from the STRING database (version 11.5) [163]. Only high confidence interactions (string score > 0.700) were computed as valid interactions. Out of the 28 *Pf*AP2 members only one, PF3D7_1115500, did not have interactors meeting these requirements. For each *Pf*AP2, the number of total interactors and total interactors inside the *Pf*AP2 family were calculated. The degree and number of interactions among *Pf*AP2s were computed and compared to a random distribution by sampling the annotated proteome of *P. falciparum 3D7*. This was chosen as background to avoid annotation bias as 32.90% (1769 proteins) of the STRING database proteome lacked annotated interactors. The size of the largest connected component and the mean shortest distance were measured according to Menche *et al.* [98]. Empirical or unpaired t test (two-sided) p-values were calculated.

### Data representation and statistical analyses

Four-way intersection Venn diagrams were made with Venny 2.1. Statistical differences between *Pf*AP2s and the *P. falciparum* (isolate 3D7) reference proteome from UniProt (proteome ID UP000001450, release 2022_03, n = 5374 proteins) [164] were analysed by a two-tailed unpaired t test with GraphPad Prism 6. For protein-protein interactions, significance of the differences was assesed by Wilcox p-value or empirical p-value.

### LLPS prediction

FuzDrop is based on conformational entropy from FuzPred method [112, 113], being able to distinguish between regions promoting aggregation or LLPS, and predict regions which mediate different interactions. FuzDrop defines a residue-based “droplet-promoting probability score” (p_DP_) and a sequence “probability of spontaneous liquid-liquid phase separation score” (p_LLPS_) which allow to distinguish the role of the protein in case of phase separation. Regions predicted as promoting droplet formation have p_DP_ ≥ 0.60 for at least 10 consecutive residues. Proteins with p_LLPS_ ≥ 60 are considered droplet drivers and p_LLPS_ <60 with at least a droplet promoting region, clients [113]. Here we refer to p_LLPS_ as FuzDrop score. PSPredictor is a machine-learning powered prediction method trained on the LLPSDB [114]. Proteins which score above the 0.5 threshold are considered as proteins capable of undergoing phase separation [114]. catGRANULE evaluates the RNA-binding capacity and protein structural disorder as well as favouring the presence of Arg, Gly and Phe (amino acids enriched in proteins undergoing LLPS) and protein length [115]. A conservative approach of proteins with granule strength ≥ 1.05 is considered as their *in vivo* experimental validation. DeePhase combines knowledge-based with a machine-learning strategy to distinguish between LLPS forming and non-forming protein sequences [117]. Scores above 0.5 are considered drivers or LLPS. Enrichment in Arg/Tyr was assessed with fLPS’s single-residue bias calculation, which searches for compositional biases above expected against a dataset of domains from ASTRALSCOP [110]. PrLD prediction was performed with the PLAAC prediction method, which evaluates input sequence compositional resemblance to yeast prions and PrLDs. Sequences with COREscore > 0 were considered positives. The PrLD boundaries were established by the PRDaa parameter [165]. The presence of soft amyloid cores inside these predicted PrLDs were scanned with pWALTZ dedicated software [102].

### Phylogenetic conservation of LLPS capacities

*Plasmodium vivax*, *Plasmodium knowlesi* and *Plasmodium malariae* proteomes were obtained from UniProt (respectively: proteome ID UP000008333, release 2022_04, n = 5389 proteins; proteome ID UP000031513, release 2022_04, n = 5339 proteins; proteome ID UP000219813, release 2022_04, n = 5915 proteins). Orthologues for the 28 *Pf*AP2 homologues were obtained from the Ortho MCL database (release 7.0, April 2025) [166], which belong to each of the assayed *Plasmodium* species and the reference proteomic arrangement coincident with each UniProt reference proteome. Additional *P. vivax* and *P. knowlesi* AP2 TF were enriched with data from Oberstaller *et* al. [167], mapped with VEuPathDB (release 68, May 2024) [168] into Gene ID, and these into UniProt accessions numbers. The final number of potential unique AP2 proteins in *P. vivax* was 30, 31 in *P. knowlesi* and 25 in *P. malariae*.

### Oligonucleotides targeting *Pf*AP2 encapsulation into ghost RBCs

Complementary DNA sequences listed in **Table S4** were used to generate the *Pf*AP2-targeting oligonucleotides. Briefly, 10 µM of each complementary oligonucleotide were mixed in tris-EDTA (TE)-Buffer, placed in a water bath at 95 °C and let to gradually cool until 25 °C to allow their annealing. The resulting dsDNA sequences were kept at −20 °C until use.

To encapsulate the oligonucleotides, ghost RBCs (gRBCs) were generated following previously published protocols [80, 119]. Briefly, human type B+ RBCs were washed twice with phosphate-buffered saline (PBS) at 200x g for 10 min at 4 °C. The remaining RBC pellet was incubated for 1 h at 4 °C under gentle stirring in cold lysis buffer (5 mM KH_2_PO_4_, 1 mM ATP, pH 7.4) containing 2 µM of each annealed oligonucleotide to encapsulate. After this time, the generated gRBCs were pelleted and half of the total volume was replaced by resealing buffer (0.5 M NaCl, 0.1 mM MgCl_2_, 10 mM ATP, 10 mM GSH) and incubated for 1 h at 37 °C under gentle stirring. Finally, samples were washed three times in Roswell Park Memorial Institute 1640 medium (RPMI) supplemented with 0.5% (w/v) Albumax II (Life Technology, Auckland, New Zealand) and 2 mM L-glutamine (complete RPMI, cRPMI), and the final oligonucleotide-encapsulating gRBCs were kept at 50% in RPMI at 4 °C until use.

The infection of gRBCs with *P. falciparum* was done by using late stage parasites purified in 70% Percoll as described previously [169, 170]. The purified parasites were added to the oligonucleotide-loaded gRBCs and incubated at 3% haematocrit in cRPMI medium. Parasites were maintained at 37 °C under an atmosphere of 5% O_2_, 5% CO_2_ and 90% N_2_ for 40 h prior to FRET experiments.

### Quantitative FRET analysis

Image acquisition and FRET efficiency by acceptor photobleaching measurements [171] were performed using a Leica TCS SP5 confocal fluorescence microscope. Before acquisition, parasites were placed in an 8 well chamber µ-slide (Ibidi) previously coated with 50 mg/mL concanavalin A, as described previously [172]. Alexa 488 and Alexa 546 were chosen as the donor and acceptor fluorophores, respectively. All samples were imaged with a Leica APO CS2 63.0 x 1.40 oil immersion lens. FRET was resolved from the increase of the donor fluorescence in the bleached region of interest (ROI) which was maintained throughout the whole experiment. Data analysis was performed in the Leica software (LASX), and the energy transfer efficiency (E) was calculated according to the following equation:

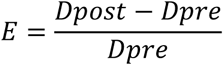

Were *Dpost* corresponds to the fluorescence intensity of the donor after photobleaching and *Dpre* is the fluorescence intensity of the donor before photobleaching. As a control, non-bleached areas were also analysed for FRET. All images were acquired by simultaneous excitation with 488 nm and 561 nm laser lines using an argon ion and a diode-pumped solid-state laser, respectively. Fluorescence emission was collected in the ranges of 498-530 nm for Alexa 488 and 567-628 nm for Alexa 546. Photobleaching was obtained by scanning over 10 vertical Z sections, with the 561 nm excitation laser line at 80% of its power intensity.

## Abbreviations

3D: three-dimensional
ACDC: AP2-coincident C-terminal domain
AP2: apetala2
FRET: Förster resonance energy transfer
gRBCs: ghost RBCs
IDP/R: intrinsically disordered protein/region
LCC: low compositional complexity
LLPS: liquid-liquid phase separation
LyRET: lyse-reseal erythrocytes for transfection
PB: photobleached
*Pf*AP2: *Plasmodium falciparum* AP2 family TF
pgRBCs: parasitized gRBCs
pI: isoelectric point
pLDDT: predicted Local Distance Difference Test
PPI: protein-protein interaction
PrD: prion domain
PrLD: prion-like domain
RBC: red blood cell
pRBC: parasitized RBC
ROI: region of interest
TF: transcription factor

## Acknowledgements

We would like to thank the Nanomalaria laboratory for helpful insights during the project. We thank the Advanced Optical Microscopy Unit-Clinic Campus from the Scientific and Technological Centres at the University of Barcelona for their support and advice with confocal microscopy.

## Ethical issues

The human blood used for *P. falciparum* cultures was commercially obtained from the *Banc de Sang i Teixits* (www.bancsang.net). Blood was not specifically collected for this research; the purchased units had been discarded for transfusion, usually because of an excess of blood relative to anticoagulant solution. Prior to their use, blood units underwent the analytical checks specified in the current legislation. Before being delivered to us, unit data were anonymized and irreversibly dissociated, and any identification tag or label had been removed in order to guarantee the non-identification of the donor. No personal data were or will be supplied, in accordance with the current Spanish *Ley Orgánica de Protección de Datos* and *Ley de Investigación Biomédica*. The blood samples will not be used for studies other than those made explicit in this research.

## Contributions

V.I. and X.F-B conceived and designed the study, V.I. and O.B. contributed to the bioinformatics analysis, Y.A-P performed the experiments, V.I. and Y.A-P analysed and designed the figures, V.I. and X.F-B wrote the final manuscript.

## Availability of data and materials

The datasets supporting the conclusions of this article are included within the article and its additional files.

## Supplementary material

**Table S1** – Disorder prediction of the *Pf*AP2 TFs. Disorder prediction according to AlphaFold 2, IUPred2a long disorder and MobiDB-lite and percentage of residues (or residue centred windows) predicted to be involved in pH-dependent or binding-dependent disorder-to-order transition.

**Table S2** – Soft amyloid cores detected inside the PrLDs of *Pf*AP2s.

**Table S3** – FuzDrop predictions for the *P. falciparum* proteome.

**Table S4** – Complementary DNA sequences used to target *Pf*AP2s.

**Table S5** – LLPS and prion-like predictions for the identified *P. vivax*, *P. knowlesi* and *P. malariae* AP2 proteins.

**Figure S1** – *Pf*AP2 protein family structural models. Computational structural models for the *Pf*AP2s obtained from the AlphaFold database coloured following the per residue predicted local distance difference test (pLDDT). As in Figure 1, models cover the globular AP2 DNA-binding domains; regions 1600-3473, 2000-3858 and 2400-4109 are depicted for the proteins *Pf*ApiAP2 PF3D7_0420300, *Pf*AP2-HS PF3D7_1342900 and *Pf*ApiAP2 PF3D7_0613800 (unmodelled in AlphaFold database). Gene ID is shown above the protein model, and the corresponding UniProt Accession number and preferred name are displayed below.

**Figure S2** – *Pf*AP2-O5 recombinant expression in *E. coli*. Induction of His(6×)-*Pf*Ap2O5. The gene sequence encoding *Pf*AP2-O5 was codon-optimized, chemically synthesized (GenScript, The Netherlands), and cloned into the pET28a(+) expression vector, introducing an N-terminal 6×His tag. Recombinant His(6×)-*Pf*Ap2O5 protein induction was carried out in *E. coli* C43 (DE3) cells for 16 hours at 16 °C in Terrific Broth Medium. Following induction, cells were lysed by sonication in His-binding buffer (20 mM sodium phosphate, 500 mM NaCl, 20 mM imidazole, pH 7.4) supplemented with 1 mM PEFABLOC SC (Roche, Germany), 0.2 mg/mL lysozyme, 20 µg/mL DNase I, and 1 mM MgCl_2_. The soluble fraction was separated from insoluble proteins by centrifugation at 17,000 × g for 30 minutes. Soluble and insoluble fractions were then separated by 12% sodium dodecyl sulphate-polyacrylamide gel electrophoresis (SDS-PAGE) and either stained with Coomassie Blue or transferred to a nitrocellulose membrane (Bio-Rad, USA). After transfer, membranes were blocked overnight at 4 °C with 5% skim milk dissolved in Tris Buffered Saline (TBS) containing 0.5% Tween 20. Subsequently, membranes were probed with anti-His(6×) monoclonal antibodies (Thermo Fisher, 1:10,000) for 3 hours at room temperature. After washing with TBS-Tween 20 (0.5%), membranes were incubated for 1 hour at room temperature with anti-mouse HRP-labelled secondary antibodies (Abcam, 1:10,000) and developed with ECL Prime Western blotting detection reagent (Cytiva). **A)** SDS-PAGE lanes from left to right display protein ladder, not induced (N/I) and induced (Ind) samples. **B)** Western blot of the same samples from the left panel probed with anti-histidine antibodies showing no expression in the non-induced sample and a minimal expression in the induced sample. The expected molecular weight of the *Pf*AP2-O5 construct was 112.4 kDa.

