## Supplementary figures and images for "Liquid-liquid phase separation (LLPS) as a sensing and adaptation mechanism: An evidence-based hypothesis on AP2 transcription factors in the malaria parasite"

### Figure S1

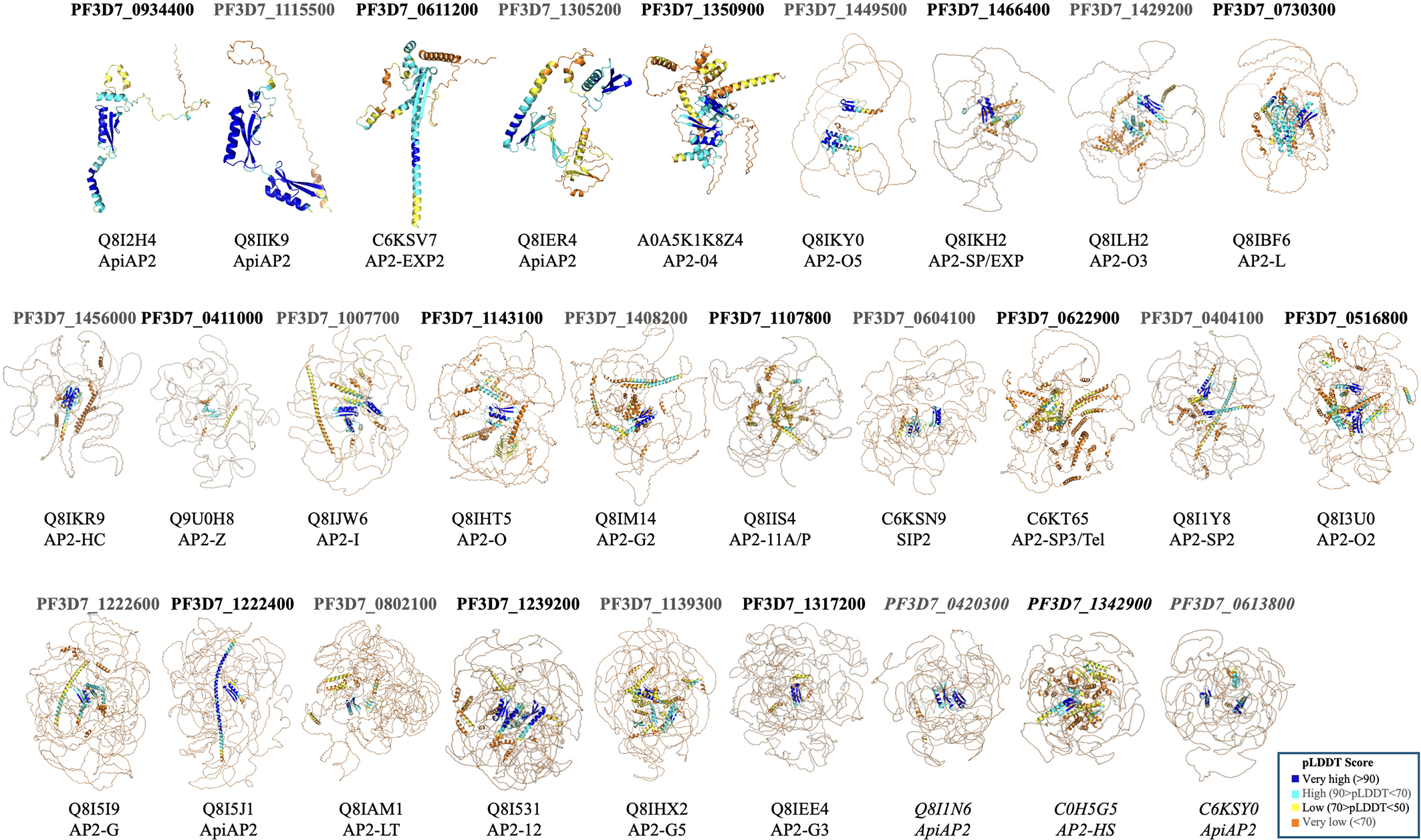

### Figure S2

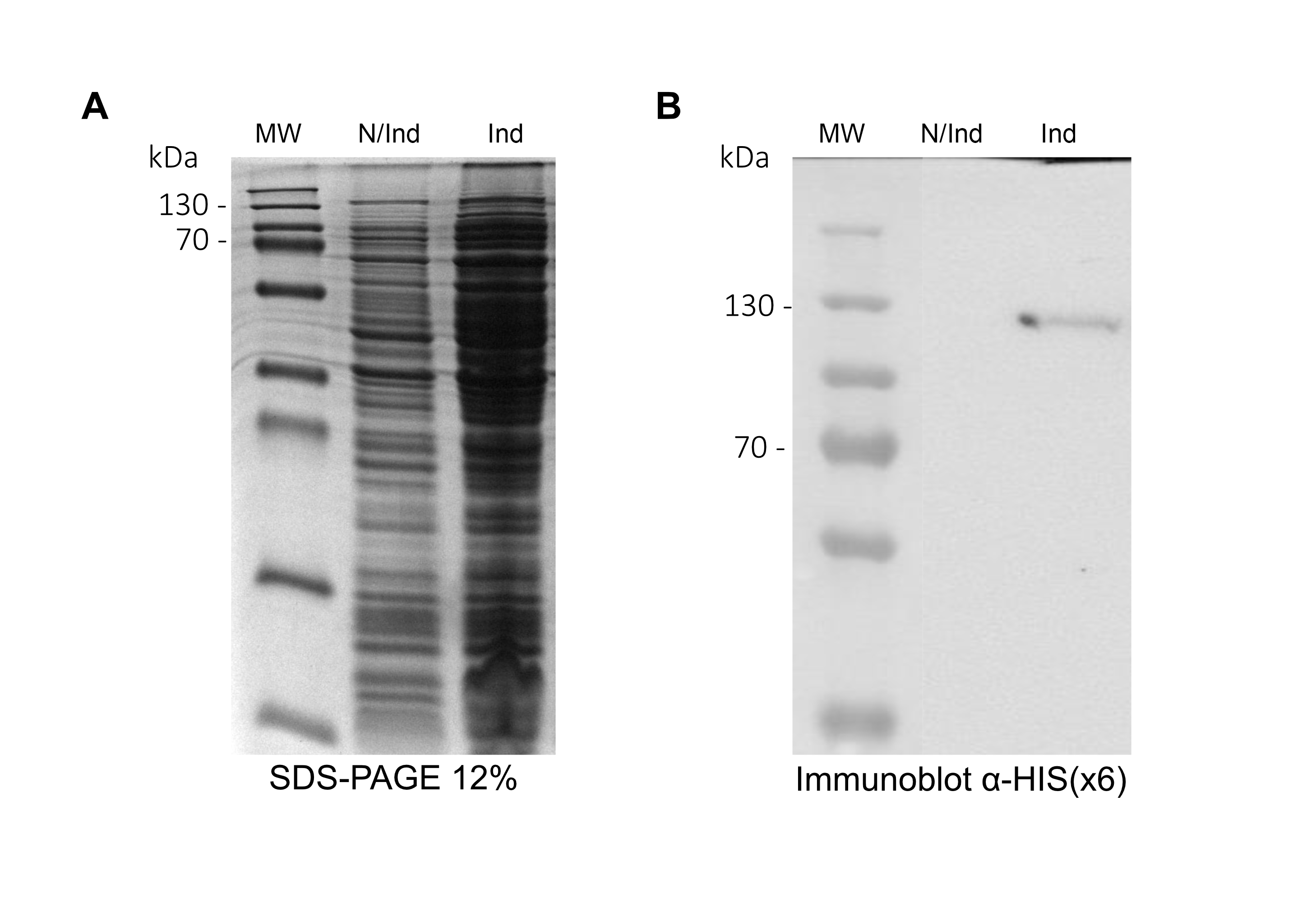
